# FakA impacts antiseptic susceptibility in *Staphylococcus aureus* and *Enterococcus faecalis*

**DOI:** 10.1101/2024.02.13.580087

**Authors:** Maria Solsona Gaya, Heather Felgate, Harsha Siani, Cynthia B Whitchurch, Mark A Webber

## Abstract

Biocides are widely used to control and prevent healthcare-associated infections and understanding how nosocomial pathogens respond to biocidal agents is key to improving infection prevention and control products and practices.

An evolution model was used to study how *Staphylococcus aureus* and *Enterococcus faecalis* responded after repeated exposure to sub-lethal concentrations of chlorhexidine digluconate (CHX) and octenidine dihydrochloride (OCT) when grown planktonically and as biofilms.

Both pathogens were able to adapt to grow at concentrations above the MIC of both biocides with planktonic lineages surviving at higher concentrations of both agents than biofilm lineages. Exposure to CHX was linked with lower biofilm biomass production in *E. faecalis* although biofilm biomass increased for *S. aureus* isolates after exposure to both agents. Evolved isolates had no major fitness deficit and only low-level changes to susceptibility to antibiotics were observed after biocide exposure.

Sequencing of biocide adapted mutants repeatedly identified mutations within *fakA* encoding a fatty acid kinase in independent lineages of *S. aureus* after exposure to both biocides in all conditions. Analogous changes were observed within the homologous gene in parallel experiments with *Enterococcus faecalis*. Further assays to study the mechanistic basis and relationship to phospholipid production showed that evolved isolates with *fakA* mutations accumulated less ethidium bromide than parent strains, exhibited altered cell envelope morphology and decreased susceptibility to daptomycin.

This data shows important pathogens can evolve limited tolerance to two common biocides but that this has collateral impacts on biofilm formation, colony morphology and fitness. FakA appears to play an important role in biocide tolerance.

## INTRODUCTION

Antimicrobial resistance (AMR) represents a major global problem (Murray *et al*., 2022) with many mechanisms of antibiotic resistance now well understood (Darby *et al*., 2023). However, much less is known about the interactions between biocides and target bacteria with mechanisms of tolerance and resistance poorly defined along with collateral impacts on antibiotic susceptibility (SCENIHR, 2010; Maillard and Pascoe, 2024). Biocide use is crucial in infection prevention and control, particularly as AMR escalates and biocides are cornerstones of strategies to control infection, especially in hospital settings (Rutala and Weber, 2004).

Biocides are widely used as disinfectants and applied to surfaces and on medical and surgical devices (Maillard, 2018; Vijayakumar and Sandle, 2019; Weber, Rutala and Sickbert-Bennett, 2019). Some biocides can also be used on skin as antiseptics with chlorhexidine digluconate (CHX) and octenidine dihydrochloride (OCT) commonly used in hospital settings to reduce bacterial carriage on skin to reduce rates of endogenous infection. Biocides must often eradicate bacteria which have formed biofilms which poses a particular challenge as biofilms are typically tolerant to antimicrobials (Percival *et al*., 2015).

*Staphylococcus aureus,* including methicillin-resistant *S. aureus* (MRSA), accounts for many healthcare-associated infections (Cassini *et al*., 2019). *S. aureus* isolates can adhere and form biofilms both within the hospital environment and on medical devices for example causing catheter-associated urinary tract infections (CAUTI) or prosthetic joint infections (PJI) (Percival *et al*., 2015). Enterococci are another frequent cause of healthcare-associated infections, especially in intensive care units (ICU) (Murray, 1990; García-Solache and Rice, 2019). *Enterococcus faecalis* (80-90% clinical isolates) and *E. faecium* (5-10% clinical strains) are both significant causes of urinary tract infections (UTI), bloodstream infections (BSI) and endocarditis (Moellering, 1992; Fiore, Van Tyne and Gilmore, 2019).

Laboratory evolution is a powerful tool to study the effects of drug exposure on bacteria and to understand the genetic basis of evolution (Poltak and Cooper, 2011; Cooper, 2018). This has recently been used to study how low levels of antibiotics impact biofilm evolution which revealed strong pressure for emergence of resistance from sub-lethal concentrations of a panel of antibiotics (Trampari *et al*., 2021). Low level exposure to some disinfectants has also been shown to be able to select for antibiotic resistance in various species (Webber *et al*., 2015; Hardy *et al*., 2018; Kampf, 2018, 2019). However, the conditions and mechanisms by which this may happen are not well understood and how biofilms may adapt to biocides is not currently clear.

To address these questions, in this study we use a laboratory evolution model to investigate the impact of prolonged exposure of *S. aureus* and *E. faecalis* to chlorhexidine and octenidine in biofilm and planktonic conditions.

## MATERIALS AND METHODS

### Bacterial strains

Two commensal isolates of *Staphylococcus aureus* (ST 188, from a nose swab) and *Enterococcus faecalis* (ST 40, from an ear swab) were selected as representatives of each species. These were chosen to represent common strain types present within a large panel of isolates recently collected from the neonatal intensive care unit (NICU) of the Norfolk and Norwich University Hospital (NNUH) where antiseptics are in regular use (Sethi *et al*., 2021). Both had been fully genome sequenced. *S. aureus* NCTC 12973 was used as a control in susceptibility testing.

### Antimicrobial agents

CHX (Sigma-Aldrich, UK) and OCT (Alfa Aesar, US) stocks were prepared in double-distilled water and dimethyl sulfoxide (DMSO) respectively. Antibiotics were purchased from Sigma-Aldrich, UK.

### *In-vitro* bead-based evolution model

An *in vitro* bead-based biofilm evolution model (Poltak and Cooper, 2011) was adapted to study the effects of exposure to two common biocides against *Staphylococcus aureus* and *Enterococcus faecalis* by exposing the isolates to escalating concentrations of CHX and OCT (for an overview see **Supplementary figure 1**). Populations were established as biofilm and planktonic lineages as in our recent work (Trampari *et al*., 2021) and studied in parallel. Four independent lineages (A, B, C, D) were established in each condition and repeatedly exposed for 72-hour windows to each test condition before being passaged. After each passage, productivity of each population, expressed as CFU/mL (for planktonic) or CFU/mm^2^ (for biofilm) were measured. The average number of generations was calculated as previously described (Poltak and Cooper, 2011) by multiplying the number of passages by log2 of the relevant dilution factor.

**Figure 1.**
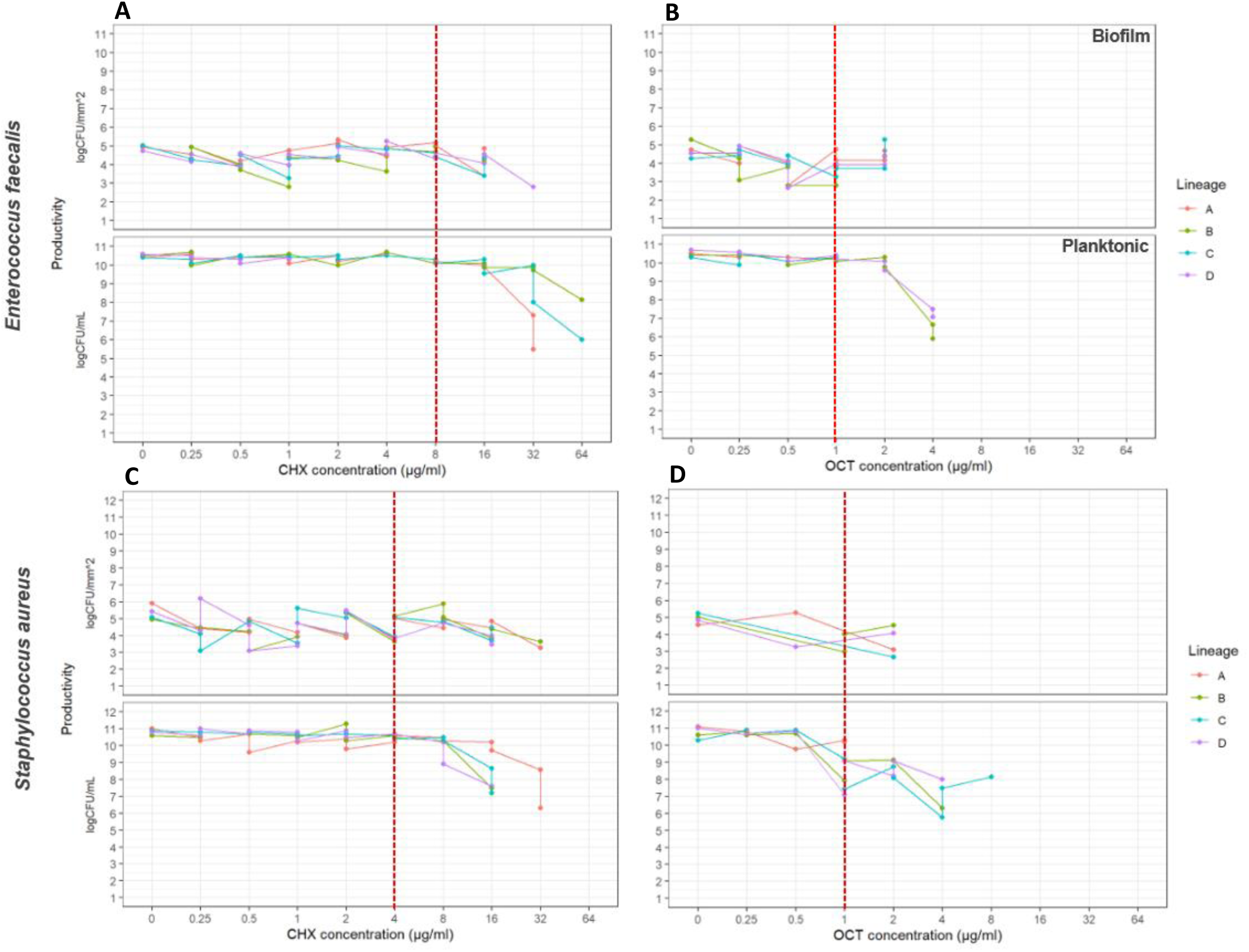
Survival of *E. faecalis* (top panels) *and S. aureus* (bottom panels) after growth in increasing concentrations of chlorhexidine (CHX) and octenidine (OCT). Productivity (expressed as CFU/mm^2^ for the biofilm and CFU/mL for the planktonic) is plotted against the biocide concentrations for both conditions (biofilm and planktonic). The MIC of the WT is shown in a red dotted line in each panel. Panel A. *E. faecalis* isolates exposed to CHX. **Panel B**. *E. faecalis* isolates exposed to OCT. Panel C. *S. aureus* isolates exposed to CHX. Panel D. *S. aureus* isolates exposed to OCT.

Biofilms were established on AISI 316 stainless steel beads (Simply Bearings Ltd, UK) in Tryptone Soy Broth (TSB) (Oxoid, Thermo Fisher Scientific, UK). To distinguish different passage points in longitudinal evolution experiments, beads of different diameters (6 mm and 3 mm) were used. The bacterial population sizes supported by the beads of different sizes were not significantly different (**Supplementary figure 2**).

**Figure 2.**
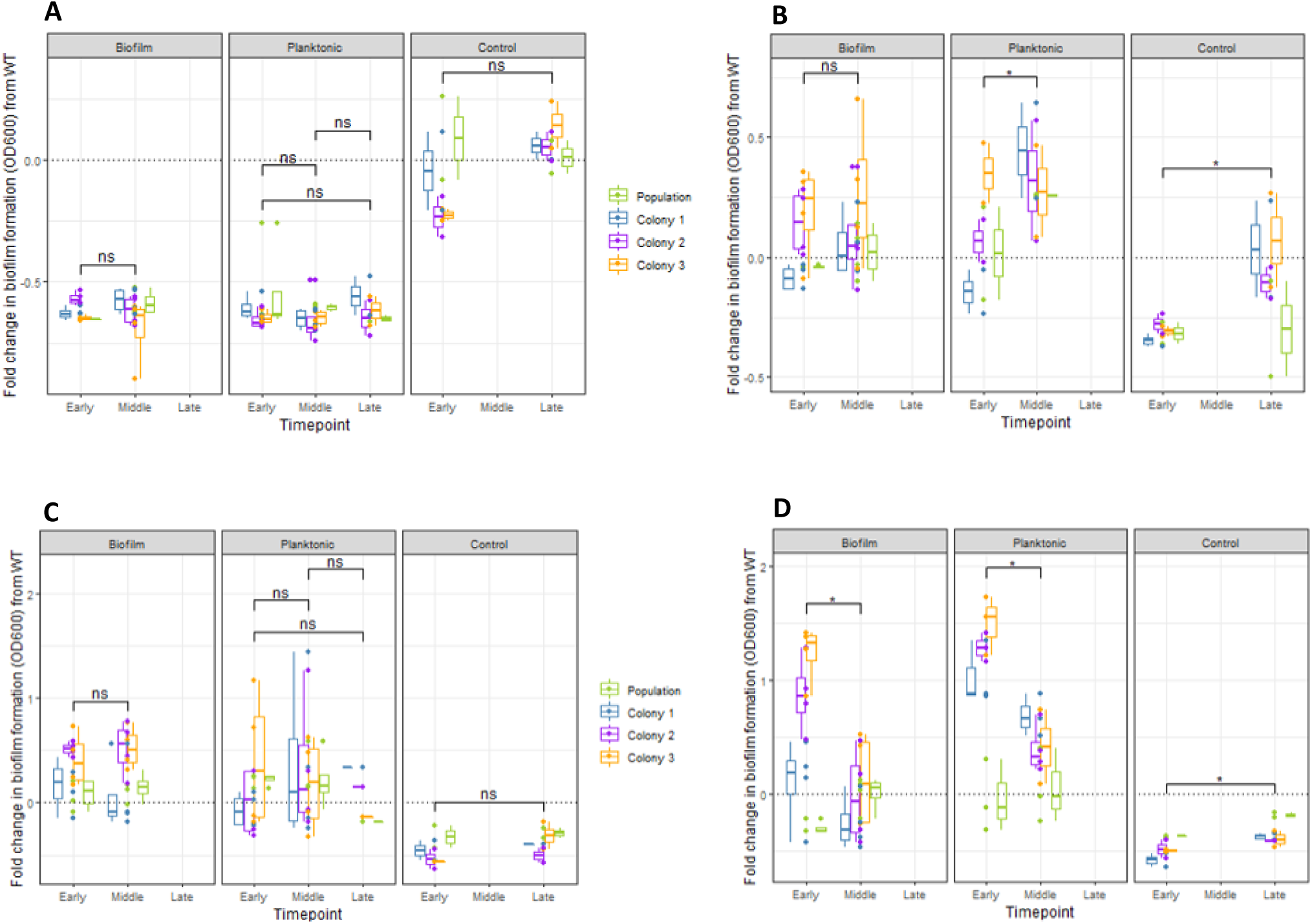
Biocide exposure had species-specific impacts on biomass production. Boxplots representing crystal violet staining with medians are shown; populations and individual isolates are represented in different colours. Fold changes in biofilm formation (OD600_nm_) from the WT are shown with the black dotted line representing the corresponding wild-type baseline. **Panel A.** *E. faecalis* isolates exposed to CHX. **Panel B.** *E. faecalis* isolates exposed to OCT. **Panel C.** *S. aureus* isolates exposed to CHX. **Panel D.** *S. aureus* isolates exposed to OCT. The Kruskal-Wallis test was used to assess differences between timepoints for each condition and the Wilcoxon-Mann-Whitney test applied as a post hoc to study differences within conditions; ns indicates non-statistically significant changes (p > 0.05) with * indicating statistically significant differences (p ≤ 0.05) within conditions according to the Wilcoxon-Mann-Whitney test.

After each passage, the beads were washed with phosphate-buffered saline (PBS) in a 24-well microtiter plate and transferred to glass tubes containing fresh TSB every 72 hours to allow bacterial regrowth. For biofilms, sterile beads were added to the glass tubes, planktonic cells were passaged by 1:100 dilutions to fresh TSB. Tubes were incubated on an orbital shaker at 40 revolutions per minute (rpm) inside a temperature-controlled incubator (22 °C). Escalating concentrations of the biocides were added to the selected conditions and increased after every two passages (from 0.25 µg/mL to 64 µg/mL). Control conditions had no biocides added. A sample of every passage was stored at −40 °C in glycerol after each passage.

### Population sampling

Populations from the start (‘early’, passage 1), ‘middle‘, and end (‘late’) of the evolution experiment were recovered, grown overnight on TSB and plated on Tryptone Soy Agar (TSA) (Oxoid, Thermo Fisher Scientific, UK). Three random colonies (C1, C2, C3) from a total of 77 populations were then selected to give 231 isolated strains. These isolates and corresponding parent populations gave a total of 308 isolates/populations which were phenotyped.

### Biofilm biomass production

Production of biofilm biomass was quantified by crystal violet staining. Overnight cultures of the desired bacterial strains were prepared in microtiter plates using 200 µL of TSB supplemented with 1 % (w/v) glucose (Sigma-Aldrich, UK) to facilitate biofilm formation (Croes *et al*., 2009; Lam *et al*., 2013; Lade *et al*., 2019). After 24 hours at 37 °C, cells were diluted (1:10000) into new microtiter plates with fresh TSB supplemented with 1 % (w/v) glucose (Sigma-Aldrich, UK). The plates were sealed with a gas permeable membrane (Thermo Fisher Scientific, UK) and incubated for 48 hours at 30 °C 200 rpm. After incubation, plates were washed with water and stained with 200 µL of 0.1 % (w/v) crystal violet (Sigma-Aldrich, UK) and incubated at room temperature for 10 minutes. Plates were washed again with water and 200 µL of 70 % (v/v) ethanol was added to each well for 10 minutes. Absorbance measurements (OD600_nm_) were determined for each well using a FLUOstar® Omega spectrophotometer (BMG Labtech, UK).

### Growth determination

Growth rates were determined as a measure of general fitness over 12 hours at 37 °C using a BMG LABTECH microplate reader (FLUOstar Omega Microplate Reader). Overnight cultures of the strains of interest were prepared in TSB and diluted 1:10000 before 200 µL were added to each well. Plates with a gas permeable seal (Thermo Fisher Scientific, UK) were inserted into the plate reader (FLUOstar Omega Microplate Reader). Growth was measured at OD600_nm_ every 30 minutes with shaking before each kinetic measurement cycle (5 flashes/well) and incubation at 37 °C.

### Antimicrobial Susceptibility Testing

Broth microdilution method was used to determine the minimum inhibitory concentration (MIC) of the tested biocides (CHX and OCT) against the control and WT samples. Agar dilution following the European Committee for Antimicrobial Susceptibility Testing (EUCAST) guidelines (EUCAST, 2024a) was used to determine the MICs of a panel of antibiotics against the 306 evolved samples. Seven antibiotics of different classes (ciprofloxacin, cefotaxime, daptomycin, erythromycin, gentamicin, penicillin, vancomycin) were tested.

Overnight cultures from each master plate were grown in Mueller-Hinton broth (MH) (Sigma-Aldrich, UK) and diluted 1:100 in MH. For daptomycin, calcium (Sigma-Aldrich, UK) was added to media to a final concentration of 50 mg/L in accordance with EUCAST guidelines. For each antibiotic, MH agar plates containing serially diluted concentrations of drug were used. Diluted cultures were spotted onto the agar (~10^4^ cells) using a multipoint inoculator (Denley Instruments Ltd, UK) and incubated at 37 °C for 24 hours. The log_2_ fold change in resistance profiles compared to the WT average was calculated for each strain.

Minimum bactericidal concentration (MBC) was determined using an agar plate assay to assess the bacterial-killing activity of the selected biocides. A total of 5 µL from the microtiter wells above the MIC (with no visible growth) were spotted on TSA plates and incubated at 37 °C for 24 hours before being examined for growth.

Minimum biofilm eradication concentration (MBEC) assays used a modified broth microdilution assay (Zaborowska *et al*., 2017). Overnight cultures were prepared in 5 mL TSB glass tubes and incubated at 37 °C. After 24 hours, cultures were diluted (1:10000) into new microtiter plates. A semi-skirted 96-well polymerase chain reaction (PCR) plate (Starlab, Milton Keynes, UK) was then allowed to sit within the wells of the microtiter tray and incubated for 24 hours (150 rpm at 37 °C) allowing a biofilm to form on the outside of the wells of the PCR plate. The microtiter tray was discarded and the outside surface of the wells of PCR plate was washed for 10 seconds in 180 µL of PBS in a microtiter tray. The PCR plate was then allowed to rest within another microtiter tray in the same format for an MIC assay (previously described). Plates were incubated at 37 °C for 18-20 hours. On day 4, the MIC plate was then discarded, and the PCR plate was placed in a fresh microtiter tray containing 180 µL of TSB which was incubated for 18-20 hours at 37 °C, when the MBEC was determined.

### DNA extraction for hybrid genome assembly

For long read sequencing DNA extractions of *Staphylococcus aureus* ST188 and selected evolved isolates were performed using the *Quick*-DNA™ High Molecular Weight (HMW) MagBead Kit (Zymo Research, UK) with the addition of 100 µL of 0.5 mg/mL lysostaphin (from *Staphylococcus staphylolyticus,* Merck, UK), and incubated for 1 hour at 37 °C. The fire Monkey kit (RevoluGen Ltd, UK) was used to extract high quality High Molecular Weight DNA from *E. faecalis* ST40. Both isolates were then sequenced on Illumina and Nanopore platforms.

For short read Illumina sequencing DNA extractions were carried out with the *Quick-*DNA Fungal/Bacterial Miniprep Kit (Zymo Research, UK), with the addition for *S. aureus* isolates, of 100 µL of 0.5 mg/mL lysostaphin and incubated for 1 hour at 37 °C.

### Whole Genome sequencing

Genomic DNA was normalized to a concentration of 5 ng/µL with 10 mM Tris-HCl. Whole-genome sequencing was performed using Illumina technology with the Flex library preparation protocol as recently described (Trampari *et al*., 2021). Long-read MinION sequencing (Oxford Nanopore Technologies) followed a protocol recently reported (Felgate *et al*., 2023).

### Bioinformatic Analysis

Genome sequencing data was stored and analysed using the Integrated Rapid Infectious Disease Analysis (IRIDA, release 19.09.2) (Matthews *et al*., 2018) and Galaxy (release 19.05) (Afgan *et al*., 2018) platforms at the Quadram Institute Bioscience. For taxonomic classification at the species level the tool Centrifuge (v0.15) (Kim *et al*., 2016) was used with FASTQ files.

From the Illumina sequencing, reads were assembled with SPAdes (v3.12.0) (Bankevich *et al*., 2012). QUAST (v5.0.2) (Gurevich *et al*., 2013) was used to check genome assembly quality and filter out bad quality samples. Assembled genomes were annotated with PROKKA (v1.14.5) (Seemann, 2014). A standard assembly and annotation pipeline included Multi-locus sequence typing assignment (MLST v0.42) (Seemann, 2016) to check sequence types and antimicrobial resistance genes which were detected using ARIBA (v2.13.2) (Hunt *et al*., 2017) with the Comprehensive Antibiotic Resistance database (CARD v3.0.1; https://card.mcmaster.ca) (McArthur *et al*., 2013). Long-read sequence data reads were assembled with Flye (v3.12.0) (Kolmogorov *et al*., 2019) and polished with Pilon (v1.20.1) (Walker *et al*., 2014). Kraken2 (v2.1.1) (Wood and Salzberg, 2014) and Bracken (v2.8) (Lu *et al*., 2017) were used for taxonomic classification and to estimate abundance, respectively.

Assembled Nanopore reads were combined with Illumina reads to create hybrid genome assemblies. Long-reads were filtered by quality with Filtlong (v0.2.0) (Wick, 2017). Illumina short reads were then mapped with Minimap2 (v2.17) (Li, 2018) and assemblies polished with two iterations of Racon (v1.3.1.1) (Vaser *et al*., 2017), one round of Medaka (v0.11.5) (https://github.com/nanoporetech/medaka). Genome completeness was estimated with CheckM (v1.0.11) (Parks *et al*., 2015) and Socru (v2.2.4) (Page and Langridge, 2019) used to validate organisation of assemblies. Assembled hybrid genomes were annotated as above.

Snippy (v4.4.3) (Seemann, 2015) was used to identify single nucleotide polymorphisms (SNPs) between reference genomes and evolved samples. Snippy-core (v4.4.3) (Seemann, 2015) combined multiple snippy outputs into a core SNP alignment. The core alignment was then run in FASTTREE (v2.1.10) (Price, Dehal and Arkin, 2009) to generate approximately-maximum-likelihood phylogenetic trees from nucleotide alignments and iTOL (v4.4.2) (Letunic and Bork, 2019) used for visualisation. Artemis (v18.2.0) (Carver *et al*., 2012) was used for genome visualisation and manual confirmation of SNPs.

### Characterisation of selected evolved isolates with FakA mutations

To further identify the role of FakA/Dak2 in biocide tolerance for both Gram-positive pathogens, we characterised six evolved isolates with SNPs in different residues of FakA. To observe how conserved the positions where changes in *S. aureus* FakA were in other related Gram-positive bacteria we retrieved homologous sequences from the NCBI database using BLAST (searching against taxID 1239, excluding staphylococci) (Altschul *et al*., 1990) before alignments of FakA sequences were performed. Sequences were uploaded into ClustalW (v1.2.4) (Sievers *et al*., 2011) for multiple sequence alignment and sequence logos were then generated with WebLogo (v3.7.12) (Schneider and Stephens, 1990; Crooks *et al*., 2004) for 100 sequences.

To measure potential changes in membrane permeability between the wild types and the six evolved isolates with *fakA* mutations, ethidium bromide accumulation was measured. Overnight cultures in MH were prepared in triplicate and incubated at 37 °C. These were diluted 1:200 in MH and incubated at 37 °C at 200 rpm until cultures reached mid log phase (OD600_nm_ = 0.2). Cultures were then exposed at room temperature for 30 minutes at ½ MIC of CHX and OCT; cells with no stress were included as controls. Tubes were centrifuged at 4000 rpm at 4 °C for 10 minutes, and pellets were resuspended in a volume of sterile PBS to normalise cultures (OD600_nm_ = 0.1). In a 96-well plate, 20 µL of ethidium bromide (to give a final concentration of 10 µM/well) and 180 µL of the cell suspensions were added to each well in triplicate. Positive control (dead cells, which were 70 % ethanol-lysed cells) and negative controls (PBS + ethidium bromide) were also included. Fluorescence (Excitation: 301 nm, Emission: 603 nm) was measured using a BMG LABTECH microplate reader (CLARIOstar®) over 6 hours with reads taken every 5 minutes. Graphs represent accumulation after 2 h 30 minutes of incubation representing a steady state.

To study cellular morphology, strains with *fakA* mutations and parent strains of both species were prepared for transmission electron microscopy (TEM). Cells were grown overnight in 5 mL glass tubes in TSB and incubated at 37 °C. Cells were centrifuged at 3000 rpm for 15 minutes (at 4 °C). Media was removed from pelleted cells which were fixed immediately with 2.5 % glutaraldehyde (Agar Scientific, UK) in 0.1 M sodium cacodylate (Agar Scientific, UK) buffer (pH 7.2) and left for 2 hours at room temperature. The fixative was replaced with 0.1 M sodium cacodylate buffer (pH 7.2). After 2 further washes with cacodylate buffer, the cell pellet was mixed with a small amount of molten aqueous 2 % low gelling temperature agarose (TypeVII, A-4018, Sigma-Aldrich, UK), which was solidified by chilling and then cut into small pieces, approximately 1 mm^3^, with a razor blade. These pieces were post-fixed in 1 % aqueous osmium tetroxide (Agar Scientific, UK) for 2 hours then dehydrated through an ethanol series (30 %, 50 %, 70 %, 80 %, 90 % and 3X 100 % dry ethanol for 15 minutes each). After the final 100% dry ethanol treatment, samples were infiltrated with LR White medium grade resin (Agar Scientific, UK) for 1 hour each in 1:1, 2:1 and 3:1 mixes of LR White resin to 100 % ethanol and finally with 100 % resin, also for 1 hour. After a 100 % resin change, the sample pieces were further infiltrated overnight on a rotator. The resin was removed and a 3^rd^ portion of 100 % LR White was added before samples were returned to the rotator for 3-4 hours. Then 4X pieces from each sample were each put into their own BEEM capsule with fresh resin and polymerised at 60 °C overnight (20-24 hours) in the oven in a fume hood. The polymerised resin was left for another 24 hours to fully cure. 90 nm thick sections were cut using a Leica UC6 ultramicrotome with a glass knife, collected on carbon coated copper grids (EM Resolutions, UK), and stained sequentially with 2 % aqueous uranyl acetate (BDH, UK) and 0.5 % aqueous lead citrate (Sigma-Aldrich, UK). Sections were examined and imaged in an FEI Talos F200C TEM at 200 kV.

### Data visualization and statistical analysis

Statistical analyses were performed using R (v4.1.2) and RStudio (v2021.09.2+381) (R Core Team, 2021; RStudio Team, 2021). Figures were produced using the package ggplot2 (v3.3.5) (Wickham, 2016).

## RESULTS

### *In-vitro* evolution in the presence of antiseptics

Susceptibility of both isolates (*S. aureus* and *E. faecalis*) to chlorhexidine digluconate (CHX) and octenidine dihydrochloride (OCT) (**Table 1**) and a panel of antibiotics (**Supplementary table 1**) were determined. The minimum inhibitory concentrations (MICs) for CHX ranged from 4-8 µg/mL and the MIC of OCT was 1 µg/mL against both isolates. Minimum bactericidal concentrations (MBC) were one dilution higher than MICs. The minimum biofilm eradication concentration (MBEC) of CHX was >64 for both isolates and 8-16 µg/mL for OCT.

**Table 1.** Minimum inhibitory concentration, minimum bactericidal concentration, and minimum biofilm eradication concentrations of the selected biocides in TSB/TSA media.

| Isolate number | MIC (µg/mL) |  | MBC (µg/mL) |  | MBEC (µg/mL) |  |
| --- | --- | --- | --- | --- | --- | --- |
|  | CHX | OCT | CHX | OCT | CHX | OCT |
| *S. aureus* NCTC 12973 | 8 | 1 | 8-16 | 1 | >64 | 16 |
| *E. faecalis* ST40 | 8 | 1 | 8-16 | 2 | >64 | 8 |
| *S. aureus* ST188 | 4-8 | 1 | 8-16 | 1 | >64 | 8-16 |

Multiple independent lineages of each strain in either biofilm or planktonic conditions were exposed to increasing concentrations of CHX or OCT (starting below the MIC before doubling every two passages) for 18 passages giving ~250 generations in biofilms and ~120 in planktonic lineages. After repeated exposure, both strains were able to adapt to grow in concentrations of biocides above the starting MIC, but growth (measured as CFU/mL for planktonic cultures and CFU/mm^2^ for biofilms recovered from beads) was compromised at the highest concentrations (**Figure 1**). Both species could adapt to grow in higher concentrations of CHX (8X the MIC) than OCT (4X the MIC). *E. faecalis* biofilms were able to adapt to escalating CHX concentrations (up to 32 μg/mL); with biofilm productivity maintained at a similar level to control conditions (~10^5^ CFU/mm^2^) until populations were exposed to the highest concentrations of CHX (**Figure 1A**). Planktonic lineages were able to grow at higher concentrations of CHX than biofilms but growth in the highest concentrations was compromised. For control lineages, growth in both planktonic and biofilm conditions remained stable across all experiments. Similar patterns of growth were observed for *S. aureus* (**Figure 1C**) as for *E. faecalis* after CHX exposure.

Both species could also adapt to escalating OCT concentrations but to lower concentrations than for CHX. Biofilms were unable to survive beyond ~4 µg/mL of OCT for *E. faecalis* (**Figure 1B**) and ~2 µg/mL for *S. aureus* (**Figure 1D**). *E. faecalis* planktonic lineages were again able to survive higher OCT concentrations than biofilms (4 µg/mL) (**Figure 1B**) and for *S. aureus* one lineage was able to survive up to 8 µg/mL of OCT (**Figure 1D**).

To determine mechanisms of antiseptic adaptation and correlate impacts on fitness and other phenotypes a panel of randomly selected isolates drawn from the start, middle and end of each experiment were characterised.

### Biofilm biomass production

Crystal violet was used to measure biofilm biomass production from isolates across the experiments. The parent *S. aureus* strain had a higher biomass production (mean OD600_nm_ = 1.374) than the *E. faecalis* strain (mean OD600_nm_ = 0.858).

Biofilm biomass production differed between isolates recovered from different conditions. *E. faecalis* mutants showed lower biofilm formation after being exposed to CHX (**Figure 2A**) than in control lineages but were able to maintain biofilm formation after exposure with OCT which increased (**Figure 2B**) when compared to the control. Biofilm formation increased after OCT exposure between early and middle time points in isolates recovered from both planktonic and biofilm conditions. In contrast, *S. aureus* isolates exposed to either biocide produced more biomass than controls (**Figure 2C and 2D**), although this reduced over time after CHX exposure (**Figure 2C**).

### Fitness measurements of the WT and evolved populations

Growth kinetics in drug free media were determined to study fitness of evolved isolates compared with controls; area under the curve values calculated and plotted relative to the WT. For both species growth in the original experiment was largely maintained until a prohibitive concentration was reached (**Figure 1**). *E. faecalis* isolates exposed to either agent recovered across the experiments were not inhibited in their ability to grow in planktonic culture compared to the parent strain and increased growth capacity slightly in line with control lineages (**Supplementary figure 3A and 3B**). *S. aureus* isolates recovered from across the time points demonstrated no significant change in growth capacity (**Supplementary figure 3C and 3D**).

**FIGURE 3.**
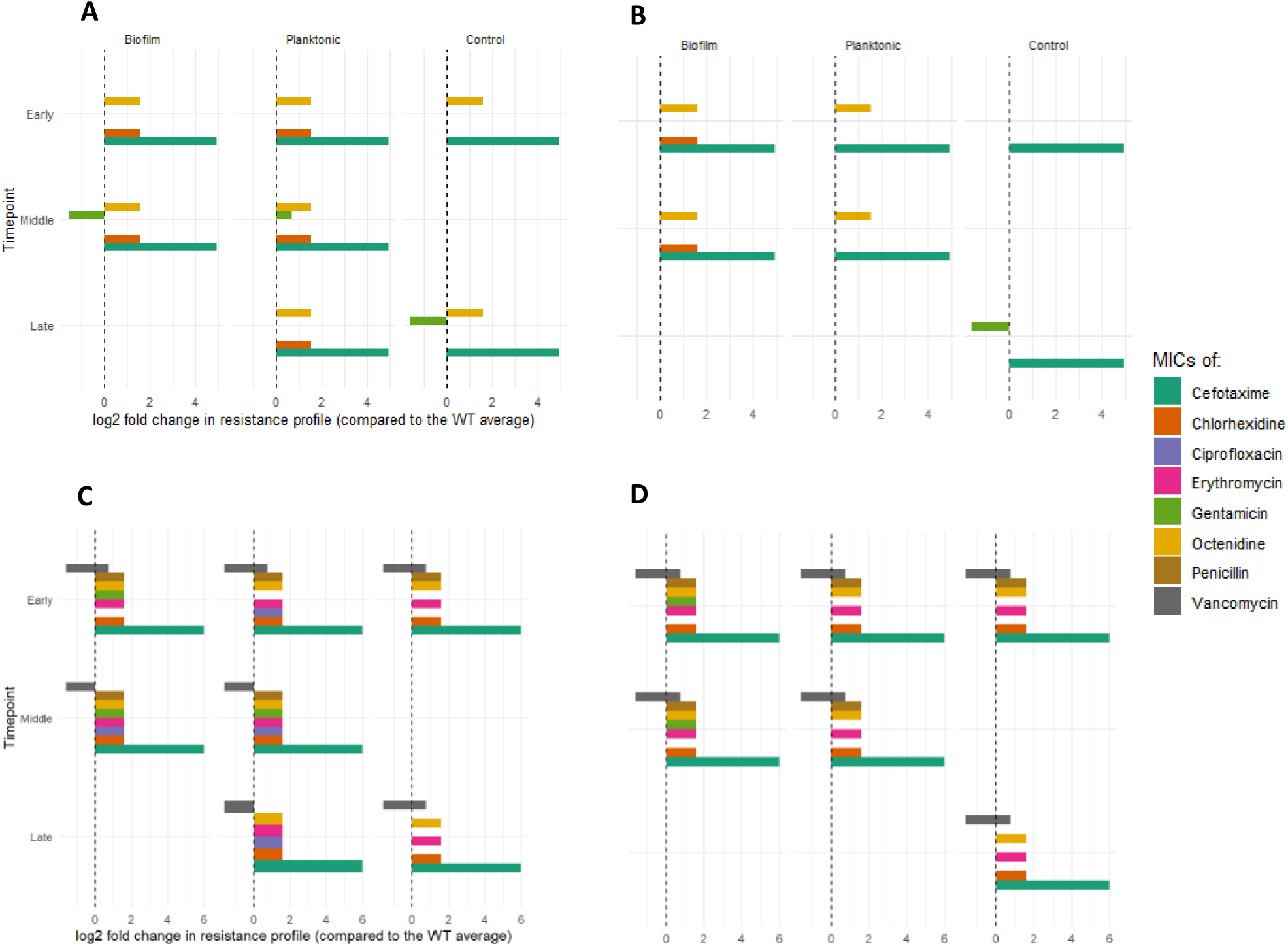
Resistance profile of *E. faecalis* and *S. aureus* evolved isolates. The log_2_ fold change in MICs (compared to the WT MIC average) was calculated for 6 different antibiotics and 2 biocides. Results for each condition (biofilm, planktonic and control) are shown. **Panel A.** *E. faecalis* CHX-exposed. **Panel B.** *E. faecalis* OCT-exposed. **Panel C.** *S. aureus* CHX-exposed. **Panel D.** *S. aureus* OCT-exposed.

### Biocide - Antibiotic cross-resistance

The MICs of both selective biocides and a panel of antibiotics were determined for the evolved isolates to identify any cross-resistance resulting from exposure to CHX or OCT. Decreased susceptibility to both antiseptics were seen in all exposed lineages although these were modest (**Figure 3**). Only low-level changes to antibiotics were observed, increases in MICs of cefotaxime were consistently observed for both *E. faecalis* (**Figure 3A and 3B**), and *S. aureus* (**Figure 3C and 3D**). For *S. aureus,* the MIC of cefotaxime against isolates surpassed the EUCAST epidemiological cut-off value (ECOFF) (4 mg/L; EUCAST, 2024b) after exposure to both drugs, however the same pattern was also seen in control lineages indicating this is a result of adaptation to the experimental conditions rather than the antiseptics. *S. aureus* isolates demonstrated a wider spectrum of other changes in susceptibility although this was usually only by 1-2 dilutions and similar changes were seen in control lineages as well.

These data show very little impact on antibiotic susceptibility occurred as a result of biocide exposure in these conditions.

### Genotyping the selected evolved isolates

After phenotyping 308 samples from across the experiments (including 77 whole populations and 231 randomly selected individual colonies), 131 samples representing final timepoints from each strain and all exposure conditions were selected for whole-genome sequencing (WGS) to determine the genetic basis behind adaptation.

Sequencing of *S. aureus* mutants identified various mutations present in the different groups and phylogenetic trees showed clear divergence according to the stressor with 5 distinct groups seen relating to different exposure to OCT or CHX (**Supplementary figure 4**). *E. faecalis* evolved isolates contained more than 600 SNPs when compared to the parental genome versus 60 SNPs found in *S. aureus* isolates. For both species, we focused on changes which independently arose in multiple lineages.

**FIGURE 4.**
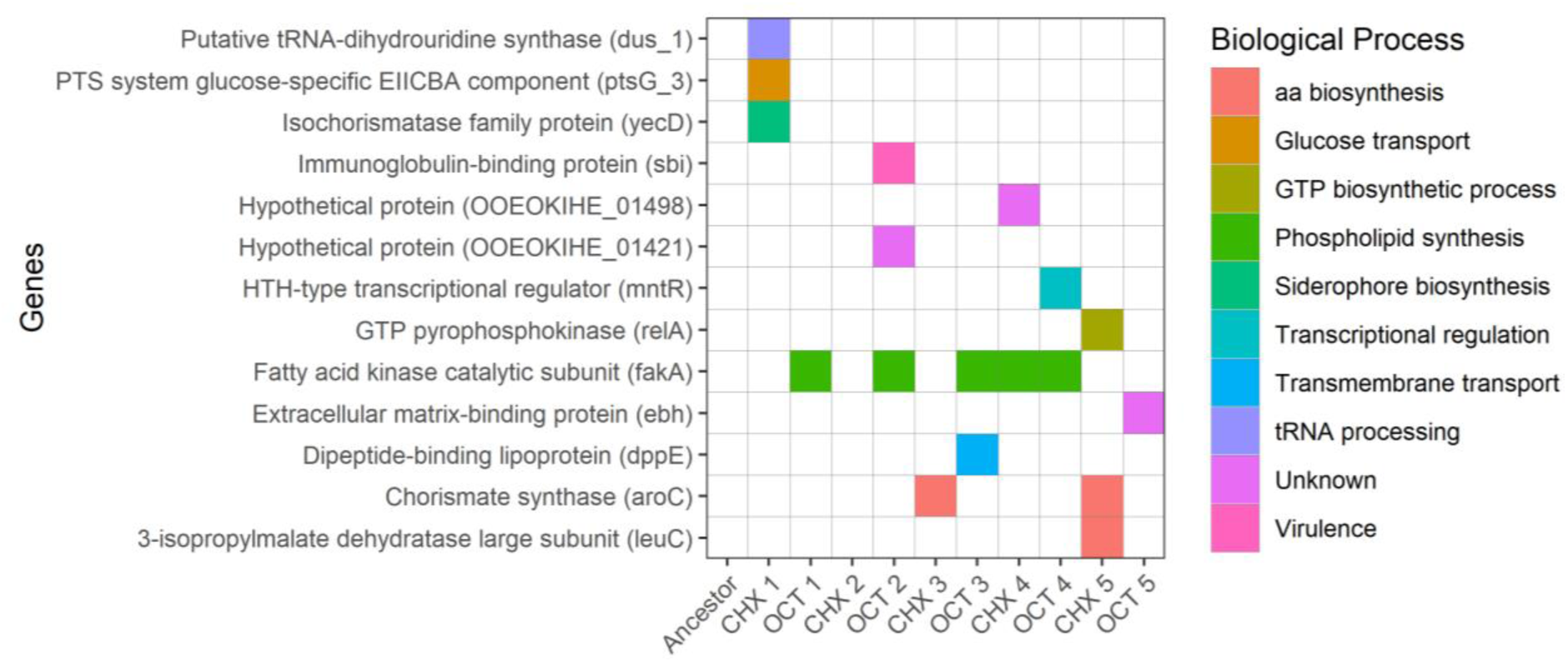
Non-synonymous SNPs present in selected evolved isolates of *Staphylococcus aureus*. Mutations in genes involved in many biological processes were found in in *S. aureus* ST188 evolved isolates stressed with CHX or OCT. Displayed are SNPs present in multiple isolates or in genes with a predicted association with antiseptic tolerance identified using Snippy-core (v4.4.3) (Seemann, 2015).

For *S. aureus*, a panel of SNPS were identified in genes involved in diverse cellular processes (**Figure 4**). Amongst these five different isolates from independent lineages exposed to either OCT or CHX carried different SNPs all within fatty acid kinase (*fakA*), which is associated with phospholipid synthesis. Two isolates presented a mutation in the chorismite synthase *aroC*.

For *E. faecalis* evolved isolates, a total of 12 SNPs were present in more than one evolved isolate. From these, 3 SNPs were found in intergenic regions and there was a mutation found in a hypothetical protein of unknown function. Of the 8 remaining SNPs, there were missense mutations in 2-hydroxyacyl-CoA dehydratase (which forms the 2-acyl-CoA product), *yitT* (required for protection against paraquat stress), *lysM*, encoding a peptidoglycan-binding domain-containing protein (with 4 SNPs in different positions of the same gene), *priA* (encoding a primosomal protein involved in DNA replication) and in a Dak2-domain containing protein where two isolates exposed to OCT presented a non-synonymous mutation within the Dak2-domain containing protein.

FakA and Dak2 are orthologues (Bose *et al*., 2014; Parsons, Broussard, *et al*., 2014) and a series of separate mutations at five different positions from 14 isolates of both species were found in the 3 different structural domains (characterised by Subramanian et al., 2022) of the fatty acid kinase which were recovered from both species in biofilm and planktonic conditions. **Figure 5** shows a graphical summary of the non-synonymous mutations in FakA/Dak2 identified in all conditions. This repeated isolation of changes within FakA indicates this is a hotspot for antiseptic adaptation.

**FIGURE 5.**
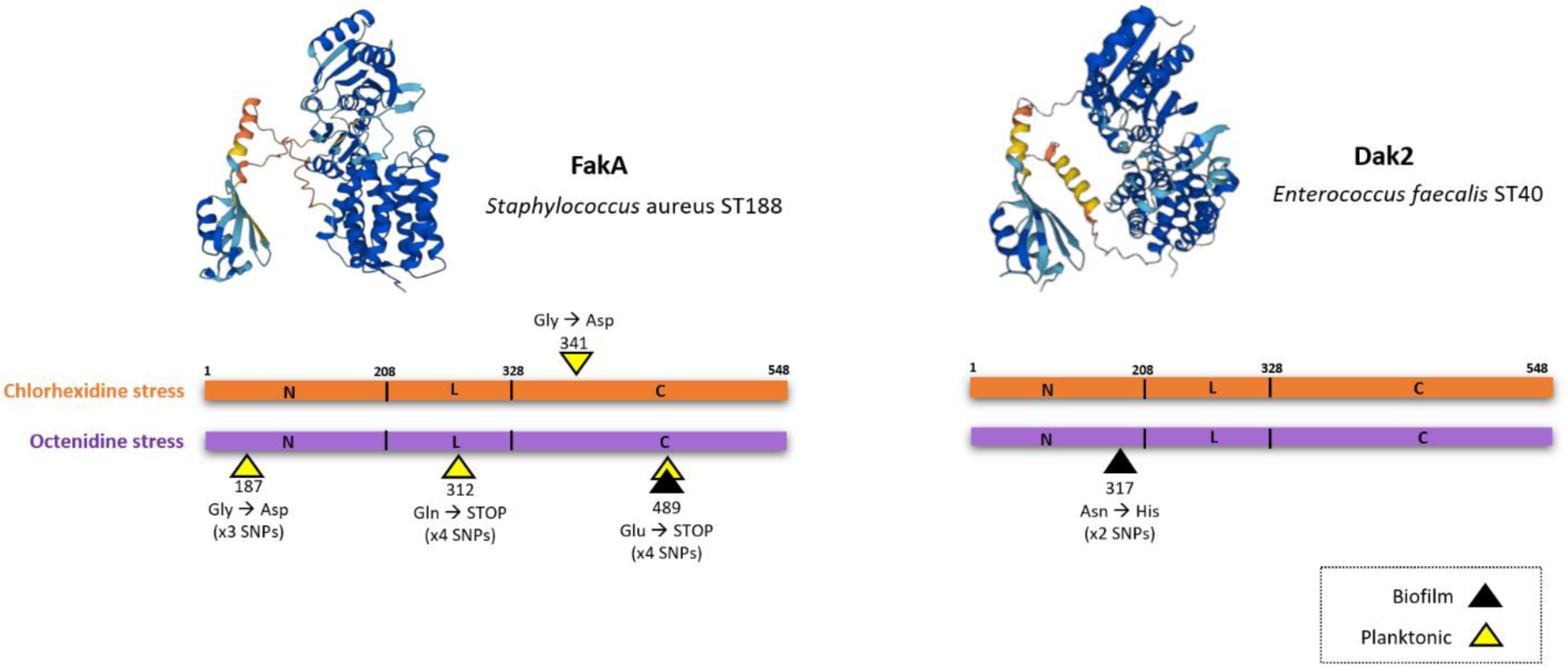
Graphical overview of SNPs present in FakA/Dak2 domains recovered from both species. Substitutions are indicated by triangles and the number of SNPs per position in the 3 main structural domains (N = amino terminal domain; L = central domain; C = carboxy domain) are shown for each stress. Predicted structure from AlphaFold DB (v.2022-11-01) by (Jumper *et al*., 2021; Varadi *et al*., 2022).

### Conservation of residues with changes in FakA

Comparison of available sequences of FakA found that residues 187, 312 and 341 (changed in mutants recovered in this study) are completely conserved across the genus *Staphylococcus* (**Supplementary figure 5**). However, residue 489 is less conserved with various substitutions apparently tolerated. Expanding analysis to other Gram-positive species found residues 187 and 312 remained completely conserved suggesting that the changes seen at these sites in this study are likely to have functional impacts on FakA.

### Phenotypic characterisation of *fakA* mutants

To assess the direct contribution of *fakA* mutation to different phenotypes, a series of mutants with different mutations within *fakA* from both species were analysed (sample IDs: 94, 176, 200, 202, 226 and 240, **Supplementary table 2**). This analysis identified small decreases in susceptibility to CHX (4/6 isolates) or OCT (6/6 isolates) and decreased susceptibility to daptomycin (4/6 isolates) (**Supplementary figure 6**).

**FIGURE 6.**
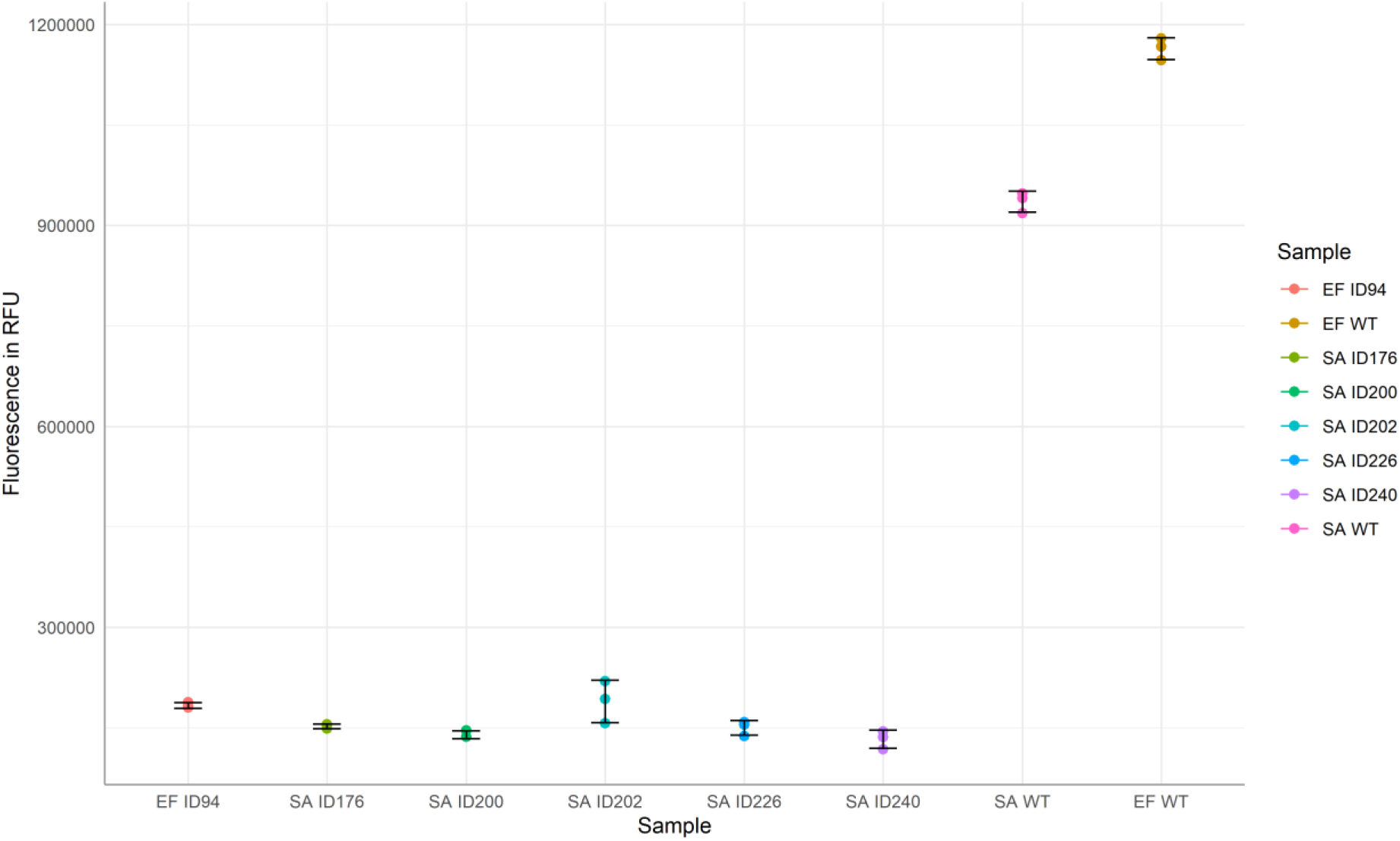
Ethidium bromide accumulation is reduced in isolates with *fakA* mutations. Ethidium bromide accumulation was measured based on fluorescence (excitation: 301nm, emission: 603 nm) of cultures over 6 hours for *S. aureus* (SA) and *E. faecalis* (EF) evolved isolates and the parent strains. Accumulation was measured after 2 h 30 minutes of incubation representing a steady state; points show 3 independent replicates per sample with error bars indicating the mean and standard deviation values of the samples.

Given the potential impact of changes to *fakA* on membrane fluidity, accumulation of ethidium bromide was also determined which showed the selected evolved *fakA* mutants all accumulated significantly less dye within the cell than the wildtypes (**Figure 6**), supporting the idea that these SNPs confer reduced membrane fluidity.

Results from TEM (**Figure 7**) showed differences in the cell envelope between the WT and evolved isolates carrying *fakA* mutations. The membranes of evolved isolates consistently showed invaginations (mesosomes) at a higher frequency than WT cells with more mesosomes per cell also being observed. In *S. aureus*, 75% of *fakA* mutants recovered after CHX treatment (and 53% after OCT treatment) showed mesosomes compared to 40% of wild-type cells. In an *E. faecalis* isolate recovered after OCT exposure with a *fakA* mutation (sample ID: 94), 75% of cells had mesosomes, whereas no wild-type cells had any mesosomes visible. The susceptibility, accumulation and microscopy results are consistent with a change to membrane composition being present in *fakA* mutants.

**FIGURE 7.**
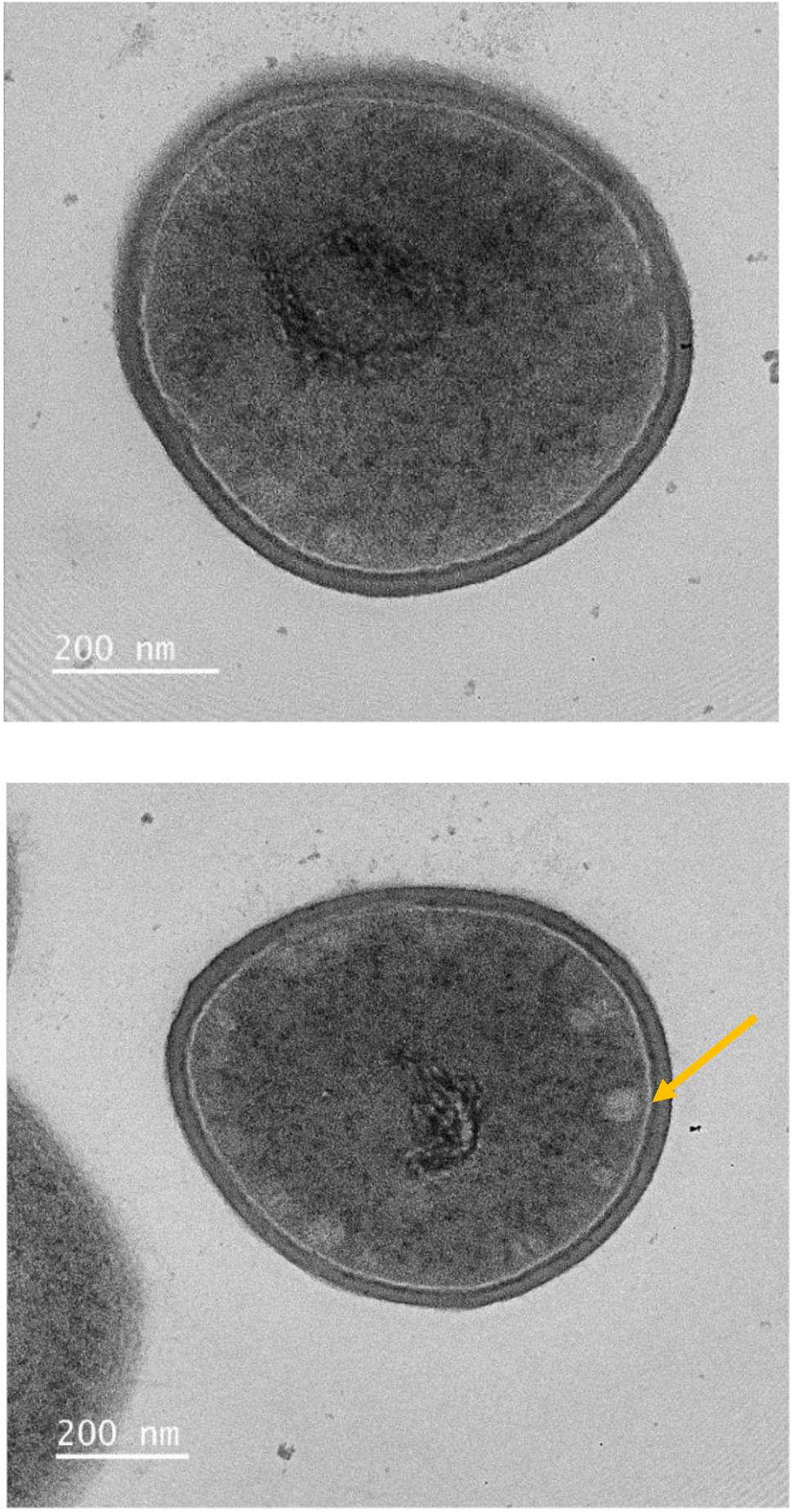
FakA mutants exhibit increased mesosome formation. Transmission electron microscopy (TEM) shows representative images of *S. aureus* WT (top) and a *fakA* mutant (SA ID176, bottom). Mesosome indicated with the yellow arrow.

## DISCUSSION

The evolution experiments showed that both species were able to adapt to grow in concentrations of CHX and OCT beyond the WT MIC and this occurred in both planktonic and biofilm conditions. However, there were collateral impacts on fitness seen in populations able to grow in escalating concentrations of the biocides; no mutants were able to adapt to grow beyond relatively low concentrations of both agents. Although biofilms are often intrinsically more tolerant of many antimicrobials, the planktonic lineages exhibited an ability to adapt to higher concentrations of both biocides than the biofilm lineages. This may reflect the larger effective population size present in the planktonic lineages but shows there is no inherently greater ability to adapt to these biocides when grown as a biofilm for both species tested.

There were some differences in the selective outcomes between the biocides and species, in general, lineages exposed to OCT retained higher biofilm-forming capacity than those stressed with CHX. Biofilm formation was also generally compromised more in *E. faecalis* after biocide exposure than for *S. aureus*.

Antimicrobial susceptibility testing results showed very limited changes in antimicrobial susceptibility followed exposure to either biocide and there were no significant changes between isolates recovered from planktonic or biofilm conditions. Changes to cefotaxime susceptibility were observed in both species but also occurred in control conditions and are likely to represent media adaptation and not a specific response to either biocide. The MIC of gentamicin (**Figure 3**) was increased for *S. aureus* isolates recovered after exposure to both biocides in in planktonic and biofilm conditions and these were above the ECOFF (2 mg/L; EUCAST, 2024). Other studies have also shown an increase in the MIC of gentamicin after CHX exposure in several *S. aureus* isolates (Wu *et al*., 2016; Nicolae Dopcea *et al*., 2020) suggesting a mechanistic link is feasible.

Whilst both biocides tested here showed limited selection for tolerance, in reality these are applied in formulation containing many other components that can impact product efficacy. Therefore, outcomes after exposure to different products containing CHX or OCT may vary.

Sequencing of mutants identified various mutations including in genes associated with amino acid and phospholipid synthesis. Two *S. aureus* evolved samples (CHX stress) presented a mutation in *aroC*, which is essential for aromatic amino acid biosynthesis. The most interesting results were found in genes associated with phospholipid synthesis. SNPs in *S. aureus* fatty acid kinase (*fakA*) were repeatedly isolated from independent lineages, for both conditions (biofilm and planktonic) and both biocides, strongly suggesting a role in biocide tolerance. SNPs in *E. faecalis* identified the presence of a mutation in Dak2 which is an orthologue of FakA (Parsons, Broussard, *et al*., 2014). The changes in FakA were mainly predicted to result in loss of function as a result of either introduction of stop codons or alteration of completely conserved residues within FakA.

FakA has recently been shown to be necessary for the incorporation of exogenous fatty acids into the lipid membrane in a pathway known as type II fatty-acid biosynthesis (FASII) bypass (Frank *et al*., 2020). Instead of using *de novo* synthesis, *S. aureus* and other Gram-positive pathogens can incorporate exogenous fatty acids and produce phosphorylated fatty acids using this pathway. An analogous system is also present in Gram-negative bacteria (Parsons, Frank, *et al*., 2014). Deletion of FakA has been shown to result in an increase in the fraction of membrane lipids containing longer acyl chains which leads to less membrane fluidity (DeMars, Singh and Bose, 2020). FakA has also been suggested to have a negative impact on biofilm formation (Sabirova *et al*., 2015) although we saw no consistent impact across the mutants in the study.

FakA homologs are present in different bacteria (e.g. *Bacillus subtilis*, *Lactobacillus johnsoni*, *Streptococcus pneumoniae,* etc.), and we found some totally conserved residues in all homologues. These include residues 187 and 312 where we saw substitutions whereas other residues where we observed changes were less conserved (341 and 489).

CHX and OCT are positively charged and interact with negatively charged phospholipids, causing membrane disruption, lysis, and cell death, although recent studies have elucidated differences in their mode of action (Rzycki *et al*., 2021; Malanovic *et al*., 2022; Waller *et al*., 2023). The repeated isolation of *fakA* mutants after biocide exposure strongly suggests a role for *fakA* in disinfectant tolerance and we hypothesised that loss of FakA function would result in altered membrane composition and as a result increased tolerance to CHX and OCT. Ethidium bromide accumulation assays showed significantly lower accumulation of this intercalating dye in the evolved isolates than in the WT. As well as this, the MIC of daptomycin, another membrane targeting antibiotic (Alder, 2005), was increased in *fakA* mutants (**Figure 6**), which is consistent with membrane alterations. In addition, TEM revealed differences in the cell wall between the WT and the evolved isolates showing increased frequency of invaginations in the plasmid membrane (mesosomes) in evolved isolates with *fakA* mutation.

Although mesosomes were regarded as artefacts of chemical fixation process in electron microscopy (Silva *et al*., 1976), there are observations that demonstrate mesosomes are a biological phenomenon in antibiotic-treated *S. aureus* (Raj *et al*., 2007). These data are again consistent with our prediction that *fakA* impacts membrane composition and as a result susceptibility to agents targeting the membrane.

## CONCLUSIONS

Mutants with increased tolerance to cationic biocides in *S. aureus* and *E. faecalis* can be selected in biofilm and planktonic conditions and biofilms are not predisposed to developing tolerance. Adaptation was relatively limited with small changes in susceptibility and little evidence of antibiotic cross-resistance. FakA appears to be a novel mediator of biocide sensitivity potentially acting by causing changes in membrane permeability.

## DATA AVAILABILITY

Genome data for the two parent isolates and the mutants have been uploaded to the National Center for Biotechnology Information (NBCI) Sequence Read Archive (SRA) at https://www.ncbi.nlm.nih.gov/sra (Accession: PRJNA1075503).

## ACKNOWLEDGEMENTS

We would like to thank the QIB sequencing team, the QIB bioimaging facilities and the QIB bioinformatics team for all their support.

## AUTHOR CONTRIBUTIONS

MSG designed and performed experiments, analysed data, and wrote the paper. HF designed the experiments, analysed data, and wrote the paper. HS helped design the study. CBW designed the experiments, analysed data, and wrote the paper. MAW designed the experiments, analysed data, and wrote the paper.

## SUPPLEMENTARY MATERIAL

**Supplementary table 1.** Minimum inhibitory concentrations (MIC) of 7 different antibiotics against *E. faecalis* ST40 and *S. aureus* ST188 determined in Mueller-Hinton (MH).

| Antimicrobial | MIC (µg/mL) |  |
| --- | --- | --- |
|  | *E. faecalis* ST40 | *S. aureus* ST188 |
| Cefotaxime | 2 | 1 |
| Ciprofloxacin | 0.25 | 0.25 |
| Daptomycin | 1 | 0.5 |
| Erythromycin | 2 | 1 |
| Gentamicin | 4-8 | 4 |
| Penicillin | 1 | 1 |
| Vancomycin | 1 | 0.5-1 |

**Supplementary table 2.** Selected *S. aureus* (SA) and *E. faecalis* (EF) evolved isolates with mutations in *fakA*.

| Species | Amino acid Residue | Sample ID | Biocide | Condition | Origin | Lineage |
| --- | --- | --- | --- | --- | --- | --- |
| SA | 187 | 202 | OCT | Planktonic | Population | D |
| SA | 312 | 200 | OCT | Planktonic | Population | C |
| SA | 341 | 176 | CHX | Biofilm | Colony 3 | B |
| SA | 489 | 226 | OCT | Biofilm | Colony 3 | D |
| SA | 489 | 240 | OCT | Planktonic | Colony 1 | D |
| EF | 317 | 94 | OCT | Biofilm | Colony 2 | B |

**Supplementary figure 1.**
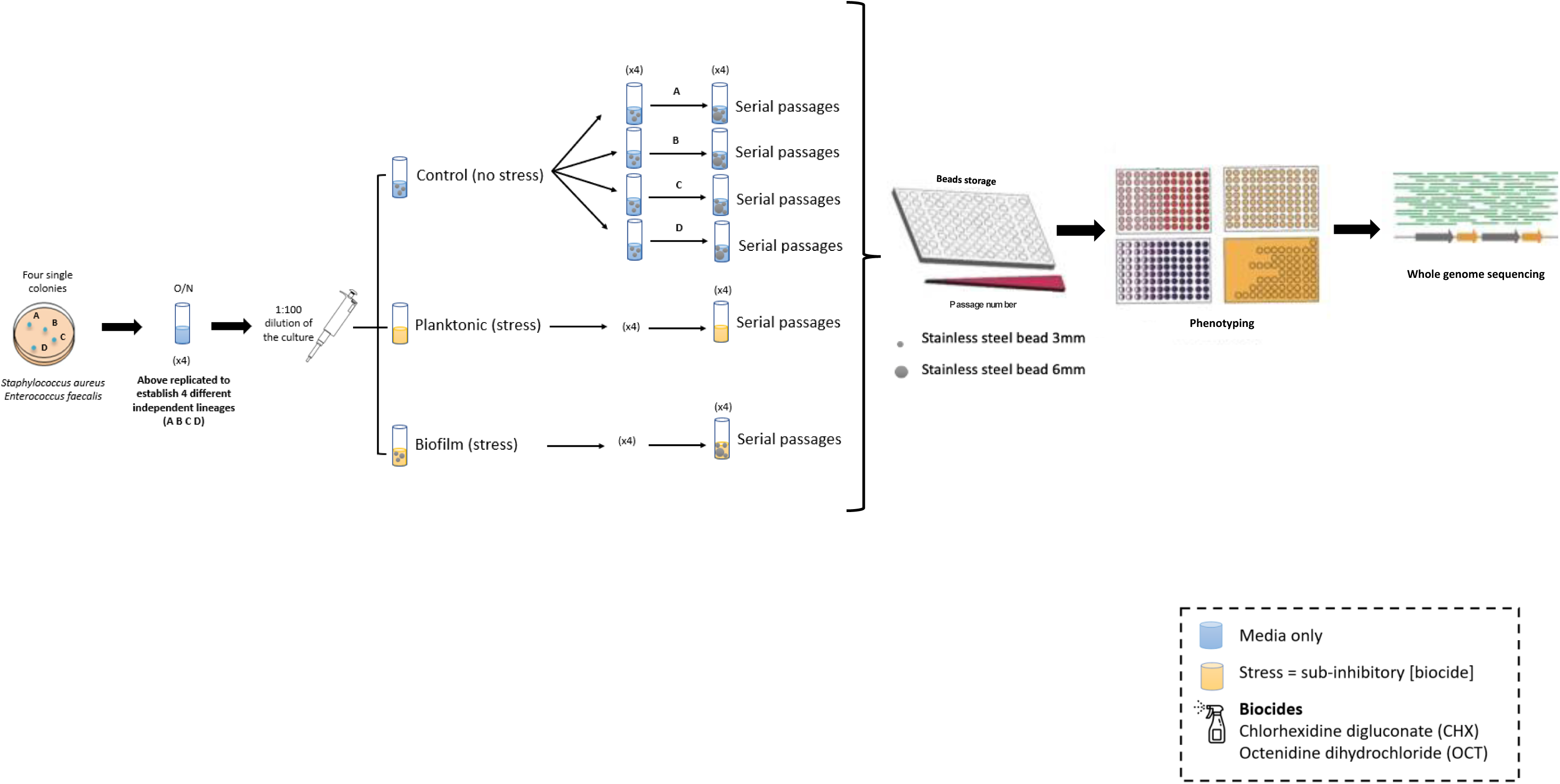
Overview of the experimental design of the *in vitro* bead biofilm evolution model.

**Supplementary figure 2.**
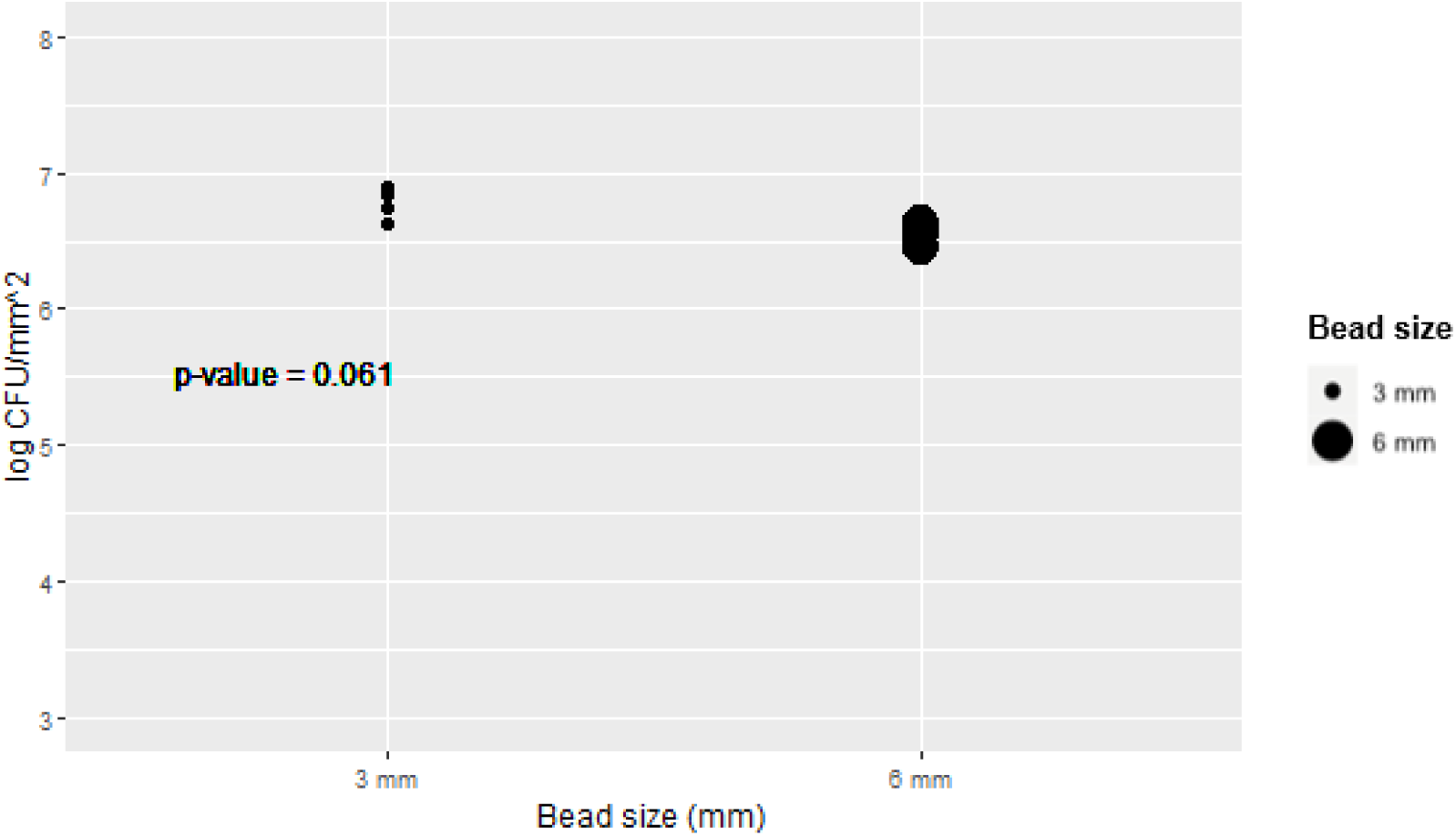
Bead size did not limit bacterial colonisation. *Staphylococcus aureus* NCTC 12973 cells recovered after growth on 3 mm and 6 mm beads were quantified (expressed as log CFU/mm^2^). Points indicate values from independent replicates from two separate experiments.

**Supplementary figure 3.**
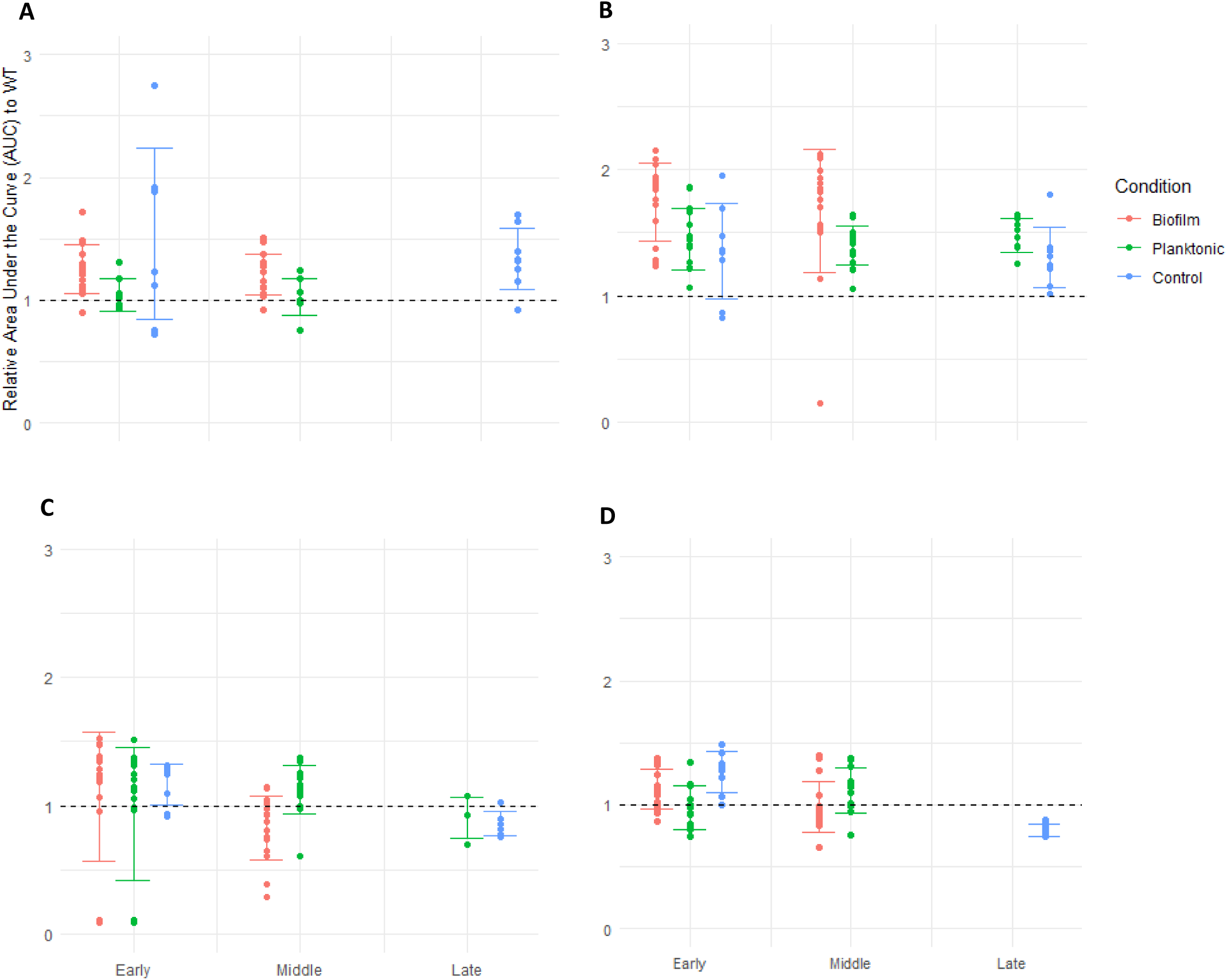
Relative growth of mutants isolated after antiseptic exposure. After growth kinetic experiments, the AUC was determined for mutants recovered from each exposure experiment relative to the parental strain (black dotted line). **Panel A:** isolates of *E. faecalis* exposed to OCT. **Panel B:** isolates of *E. faecalis* exposed to CHX. **Panel C:** isolates of *S. aureus* exposed to CHX. **Panel D:** isolates of *S. aureus* exposed to OCT. Error bars determined by the mean and standard deviation values of the samples, grouped by condition.

**Supplementary figure 4.**
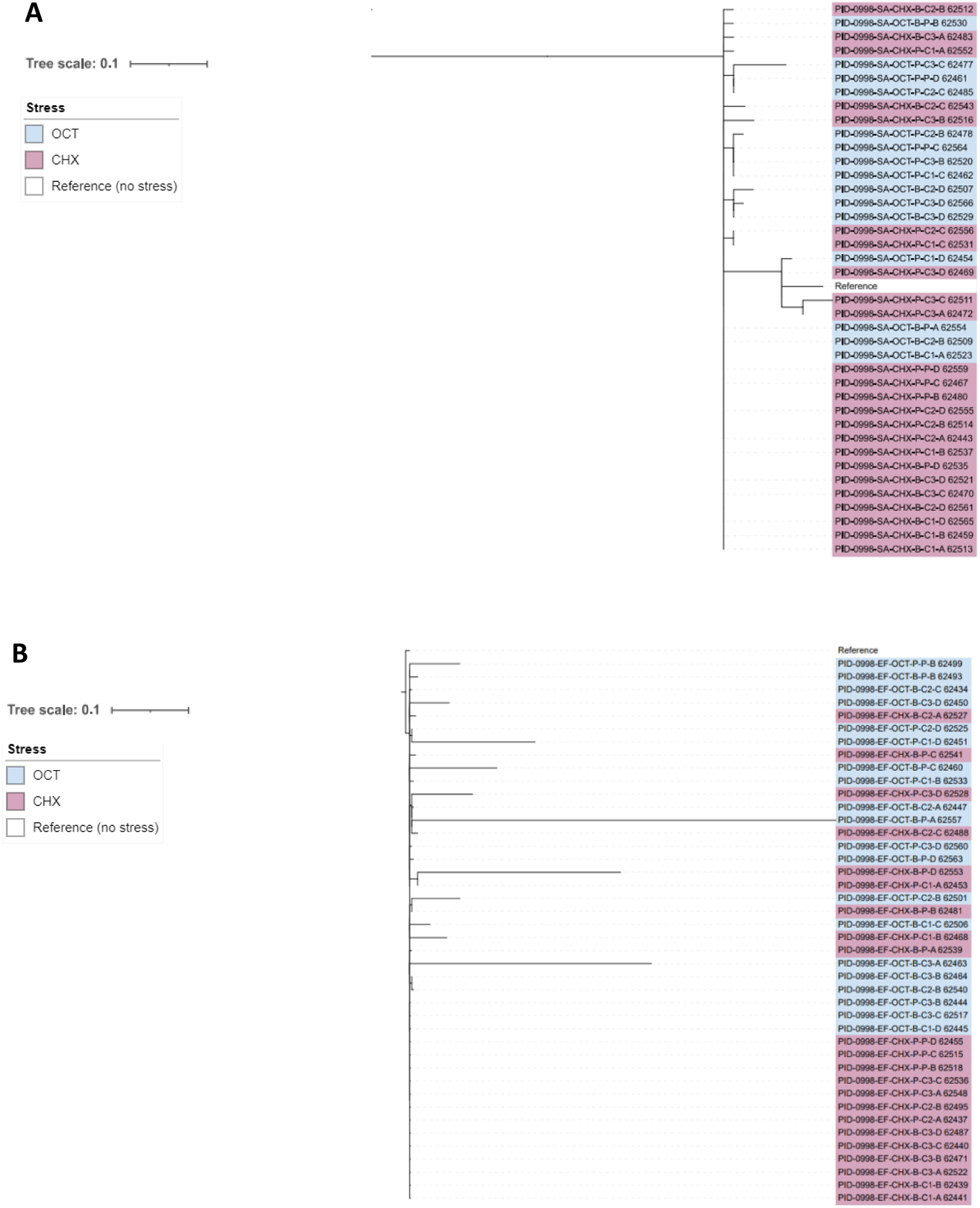
Phylogenetic trees from alignment of *S. aureus* ST188 (A) and *E. faecalis* ST40 (B) evolved isolates with the reference wildtype genomes. Approximately-maximum-likelihood phylogenetic trees from Snippy-core (v4.4.3) (Seemann, 2015) alignments generated with FASTTREE (v2.1.10) (Price, Dehal and Arkin, 2009). The phylogenetic trees were rooted using the reference strain. Results shows different groups selected by different stress conditions (CHX highlighted in pink, OCT in blue). **Legend:** Species (SA/EF), Biocide (OCT/CHX), Condition (Planktonic/Biofilm), Origin (Population/Random colony [C1, C2, C3]), Lineage (A, B, C, D).

**Supplementary figure 5.**
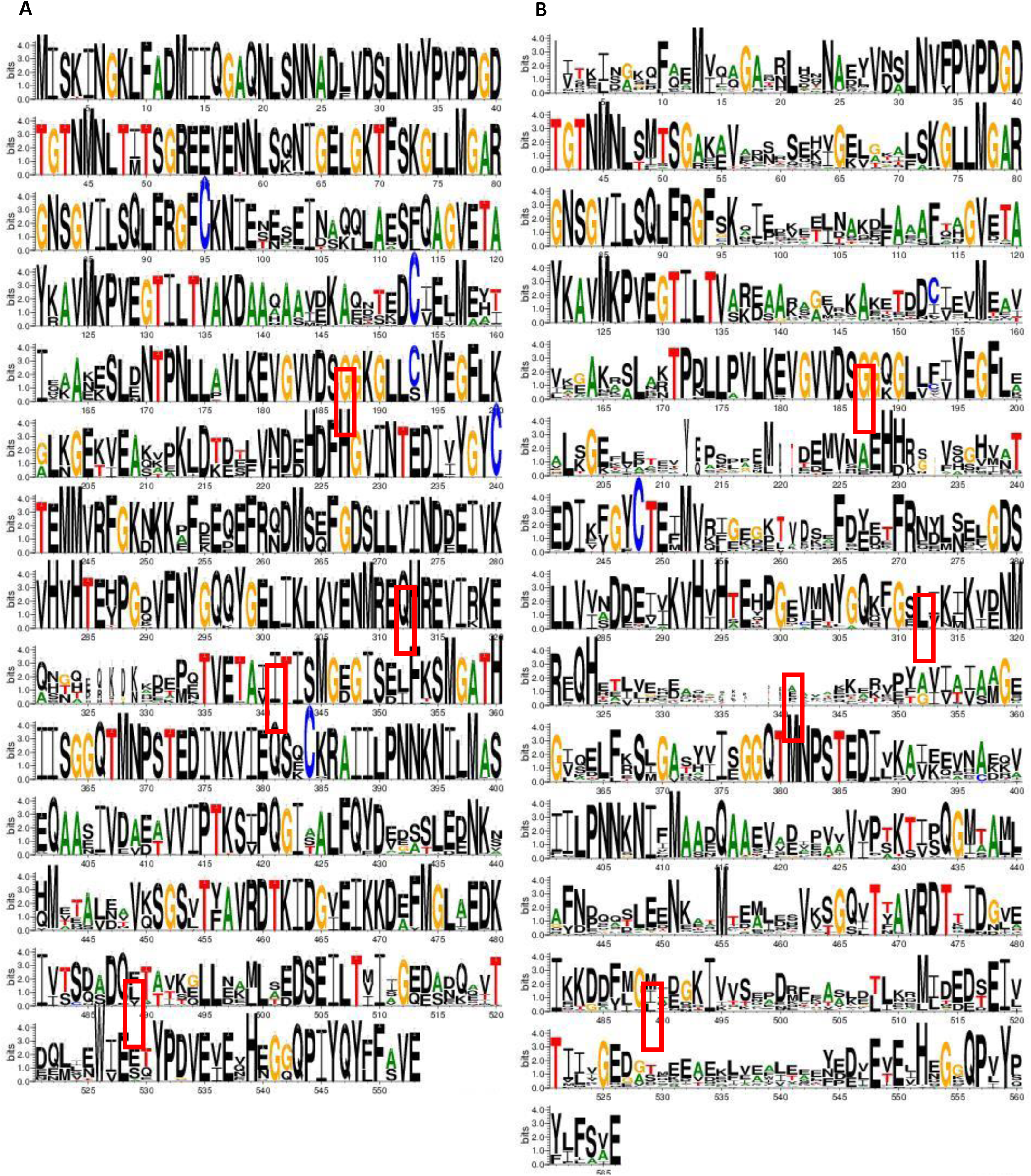
Sequence logo of FakA residues in (A) staphylococci excluding *S. aureus* and in (B) related non-staphylococcal species. The data for this logo consists of 100 sequences. Positions with SNPs in FakA (187, 312, 341, 489 aa) are highlighted in a red box.

**Supplementary figure 6.**
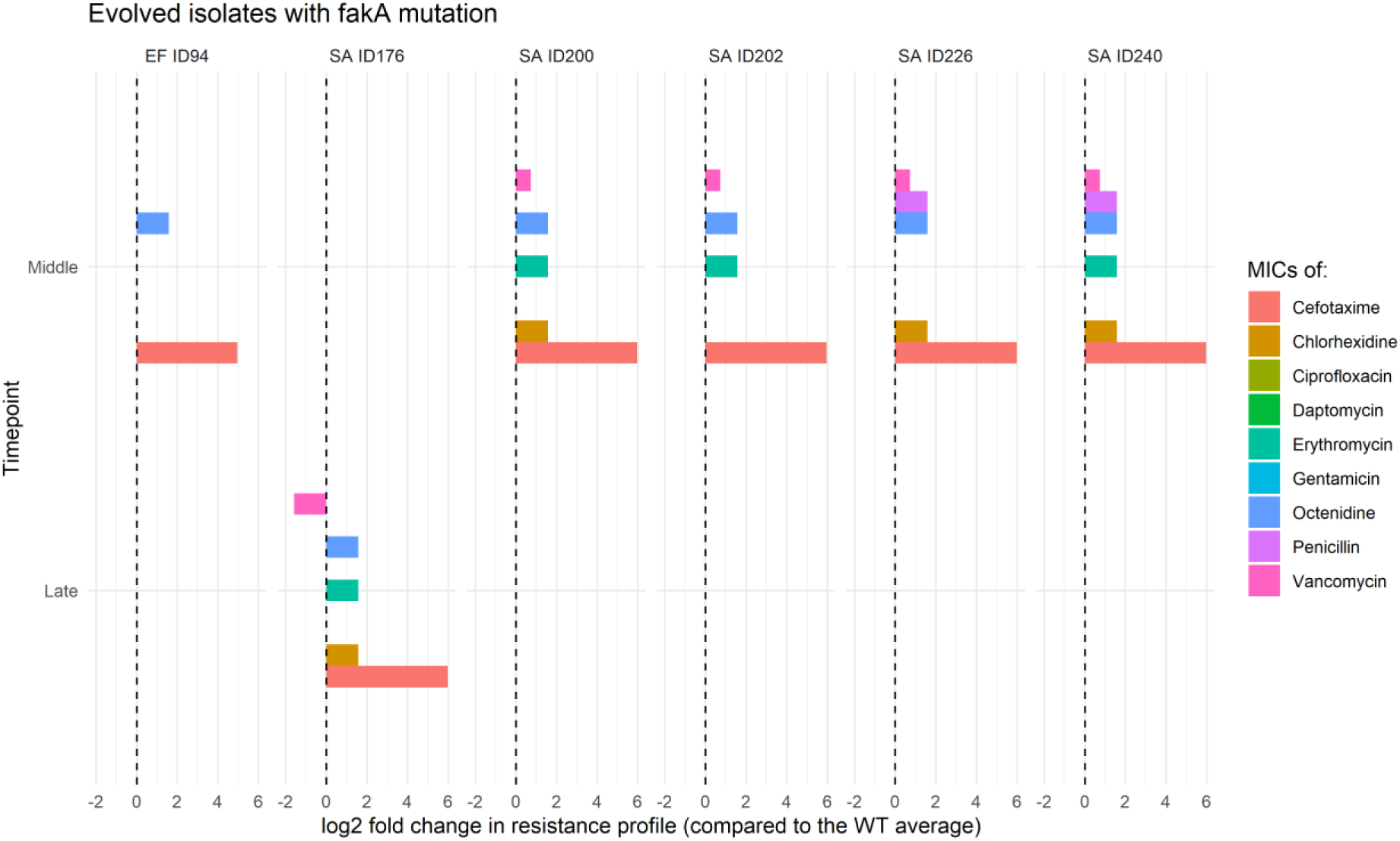
Resistance profile of selected evolved isolates with *fakA* mutation. The log_2_ fold change in MICs (compared to the WT MIC average) was performed for the 6 selected samples for 7 different antibiotics and 2 biocides. Low-level changes to various drugs observed.

